# Goal-directed action is transiently impaired in an hAPP-J20 mouse model of Alzheimer’s disease, and in aging male mice regardless of genotype

**DOI:** 10.1101/2022.07.25.501480

**Authors:** Amolika Dhungana, Serena Becchi, Jessica Leake, Gary Morris, Nesli Avgan, Bernard W. Balleine, Bryce Vissel, Laura A. Bradfield

## Abstract

Cognitive-behavioural testing in preclinical models of Alzheimer’s disease has typically been limited to visuo-spatial memory tests and has failed to capture the broad scope of deficits patients also display in goal-directed action control. The current study addresses this gap by providing the first comprehensive investigation of how goal-directed actions are affected in a transgenic mouse model of Alzheimer’s disease. Specifically, we tested outcome devaluation performance – a popular test of goal-directed action – in male and female human amyloid precursor protein (hAPP)-J20 mice. Mice were first trained to press a left and right lever for unique pellet and sucrose outcomes respectively (counterbalanced) over four days. On test, mice were fed one of the two outcomes to reduce its value via sensory specific satiety and subsequently given a choice between levers. Goal-directed action was intact for 36-week-old wildtype mice of both sexes, because they responded more on the lever associated with the still-valued outcome than that associated with the devalued outcome (i.e. Valued > Devalued). Goal-directed action was impaired (Valued = Devalued) for J20 mice of both sexes, and for 52-week-old male mice regardless of genotype. Following an additional 4 days of lever press training (i.e., 8 days lever pressing in total), outcome devaluation was intact for all mice regardless of age or genotype. Immunohistochemical analysis revealed that increased microglial expression in the dorsal CA1 region of the hippocampus was associated with poorer outcome devaluation performance on initial tests, but not with tests performed after 8 days of lever pressing. Together, these data demonstrate that goal-directed action is transiently impaired in J20 mice of both sexes and in aging male mice regardless of genotype, and that this impairment is related to neuroinflammation in the dorsal CA1 region of the hippocampus.

## Introduction

Alzheimer’s disease causes devastating impairments in memory, cognition, and behaviour (Kidd, 2008; Wenk, 2003) yet remains largely untreatable; over 99% of treatments tested in Phase 2 and Phase 3 clinical trials have failed (Anderson et al., 2017; Cummings, Morstorf & Zhong, 2014). The neuropathological features of Alzheimer’s include neuronal loss, synapse loss, increased neuroinflammation, tau hyperphosphorylation leading to the formation of neurofibrillary tangles (Delacourte, & Defossez, 1986), and amyloid-beta (Aβ) aggregation into plaques (Glenner & Wong, 1984; Trejo-Lopez, Yachnis & Prokop, 2021). Historically, it is this latter feature that has been considered its most central characteristic (Bharadwaj et al., 2009) yet numerous treatments that have successfully cleared Aβ plaques as well as produced cognitive-behavioural recovery in rodent models have failed to produce similar cognitive-behavioural recovery in humans (Cummings, Morstorf & Zhong, 2014; Hung & Fu, 2017). Better approaches to translatability are clearly needed.

One facet of rodent models that could be straightforwardly improved are the behavioural and cognitive tests used to determine how well such models replicate the cognitive-behavioural impairments of Alzheimer’s disease. According to the National Institute of Ageing (NIA), an individual will only be diagnosed with Alzheimer’s if their symptoms of memory loss and visual/spatial problems are “significant enough to impair a person’s ability to function independently”, yet rodent models routinely test only locomotor, spatial, or memory deficits (using assays such as open field, rotarod, and radial arm maze) without any regard to how these issues might translate into functional outcomes. Therefore, it was the aim of the current study to provide the first investigation of goal-directed action control in a preclinical, hAPP-J20 mouse model of Alzheimer’s disease. We hoped to provide an example of how such tests could also be administered in other transgenic mouse models and, ultimately, be used to test the capability of treatments to rescue function beyond visuo-spatial abilities, thus addressing a broader scope of Alzheimer-like deficits and increasing translatability.

Tests of goal-directed action, such as outcome devaluation, involve the integration of multiple cognitive processes including memory, motor control, and motivation. Outcome devaluation has also been translated to humans and tested in individuals with several different diseases and disorders (Morris et al., 2020; Wong et al., 2018) where they have been linked to functional outcomes (Byrne et al., 2018; Sjeords et al., 2016). For these reasons we chose to explore outcome devaluation performance using the hAPP-J20 mouse model of Alzheimer’s disease generated by Mucke et al (2000). We chose this transgenic model for two reasons. First, as this line expresses human APP (hAPP) bearing two mutations; the Swedish (K595N) and Indiana (M596L) mutation, it leads to increased Aβ plaque load, neuronal loss, and neuroinflammation relative to wildtypes. These neuropathological features have been extensively characterised in the dorsal CA1 region of the hippocampus (Wright et al., 2013; Yiu, Rashid & Josselyn, 2011) which is of interest because it is the same region we recently demonstrated to play a temporally transient role in goal-directed action (Bradfield et al., 2020), leading us to hypothesise that J20 mice might display a similar impairment. Second, J20 mice have demonstrated clear cognitive/behavioural deficits across several of the ‘traditional’ tests we have mentioned above, including increases in locomotor activity (Cheng et al., 2007), impaired spatial memory in tests such as the radial arm maze (Wright et al., 2013), and Barnes maze (Flores et al., 2021) and Morris water maze (Yoshikawa et al., 2018; Cissé et al., 2011), suggesting that these mice are behaviourally abnormal and thus an ideal initial candidate for also identifying impairments in goal-directed action.

We tested outcome devaluation in the hAPP-J20 mouse model of Alzheimer’s after different amounts of lever press training to determine exactly when such deficits occur and whether they could be overcome with additional training. In our prior study examining the role of dorsal CA1 in goal-directed action (Bradfield et al., 2020), we first trained rats to press two levers (left and right) for two unique outcomes (pellets and sucrose, counterbalanced) over 1-2 days. We then reduced the value of one of these outcomes by feeding it to satiety (Balleine & Dickinson, 1998), and subsequently gave rats a choice test during which both levers were extended but no outcomes delivered. Control animals with an intact CA1 demonstrated accurate devaluation performance by selectively responding on the lever that, during training, had earned the valued outcome and avoiding the lever associated with the devalued outcome. Because this test was conducted in extinction (i.e., no outcomes were delivered) these animals must have retrieved the lever press-outcome contingency from memory. Their intact devaluation performance was therefore evidence of goal-directed control, as it was motivated by both the current value of the outcome, as well as the contingency between the action and outcome – the two criteria of goal-directed action proposed by Dickinson and Balleine (1994). By contrast, goal-directed action was impaired for animals that received pharmacological (muscimol) or chemogenetic inactivation of the dorsal CA1 region and subsequently responded equally on the valued and devalued levers. This effect was temporally transient because when animals were tested after additional lever press training or resting in their home cages for one week, devaluation performance was intact for all rats, regardless of the status of their dorsal hippocampus.

Based on these findings, we hypothesised that J20 mice would likewise express a transient impairment in outcome devaluation performance, as a result of the damage they suffer to their dorsal CA1 region. Specifically, we expected J20 mice to be initially impaired in their outcome devaluation performance, but that they would overcome this deficit with additional lever press training. Following behavioural testing we quantified immunohistochemical markers of Aβ deposition (Amylo-Glo), and of neuroinflammation such as ionized calcium binding adaptor molecule 1 (IBA1) and glial fibrillary acidic protein (GFAP) in the dorsal CA1 region of a subset of mice, which we then compared between groups and correlated with a measure of devaluation performance from both tests, finding that increases in microglial expression in this region were associated with poorer initial goal-directed control.

## Results

### Goal-directed action is initially impaired for J20 male mice at 36-week-old and for all 52-week-old male mice regardless of genotype

Thirty-nine male mice, comprised of 4 groups: 36-weeks-old wildtypes (n = 8), 36-week-old J20 mice (n = 9), 52-week-old wildtypes (n = 8), and 52-week-old J20s (n = 14), were used for this study. Ages were taken at the beginning of behavioural training. We chose to test mice at 36-weeks because this was the earliest age at which Aβ plaque load was previously reported to significantly differentiate from wildtypes (Wright et al., 2013), and 52-weeks was chosen to determine whether the behavioural profile of these animals changed as the animals aged and neuropathological features progressed.

We have reported the data for female mice separately below because slightly different experimental parameters were used for male and female mice (see below and materials and methods for details). Regrettably, due to historically limited breeding of female mice, we were only able to age one cohort of females to 36-weeks for this study such that we did not have a 52-week-old female cohort for comparison. Current breeding protocols in our laboratory are aimed at overcoming this historic bias such that all our future studies contain equal numbers of male and female animals.

The outcome devaluation procedure used is shown in Figure 1A. As mentioned, we based the current study on the prior investigation by Bradfield et al. (2020) that was conducted in rats. However, our pilot studies revealed that alterations were necessary in order to translate this task to mice, because 1-2 days of lever press training was not sufficient for wildtype mice to demonstrate intact devaluation. Therefore, we initially trained mice to press a left and right lever (in Figure 1A these actions are referred to as A1 and A2 respectively, counterbalanced) for pellet and sucrose outcomes (in Figure 1A these outcomes are O1 and O2 respectively, counterbalanced) over 4 days on an increasing ratio schedule (see materials and methods for details). Half of the mice in each group received the left lever paired with pellets and right lever with sucrose, and the other half received the opposite arrangement (counterbalanced). Mice then received the initial devaluation test, which we hereafter refer to as the “4 Day test”, during which one of the two outcomes (O1) was pre-fed to satiety to reduce its value. Mice were subsequently given a choice between the two levers in extinction (i.e., both levers were extended but presses did not earn any outcomes). The test was repeated on the following day with the alternate outcome, O2. Animals with intact goal-directed action should respond on the lever associated with the still-valued outcome (Valued > Devalued, in Figure 1A: A1 > A.2). We expected devaluation to be intact for wildtype animals of either age, but that it would be impaired (i.e., Valued = Devalued) for J20 mice.

**Figure 1.**
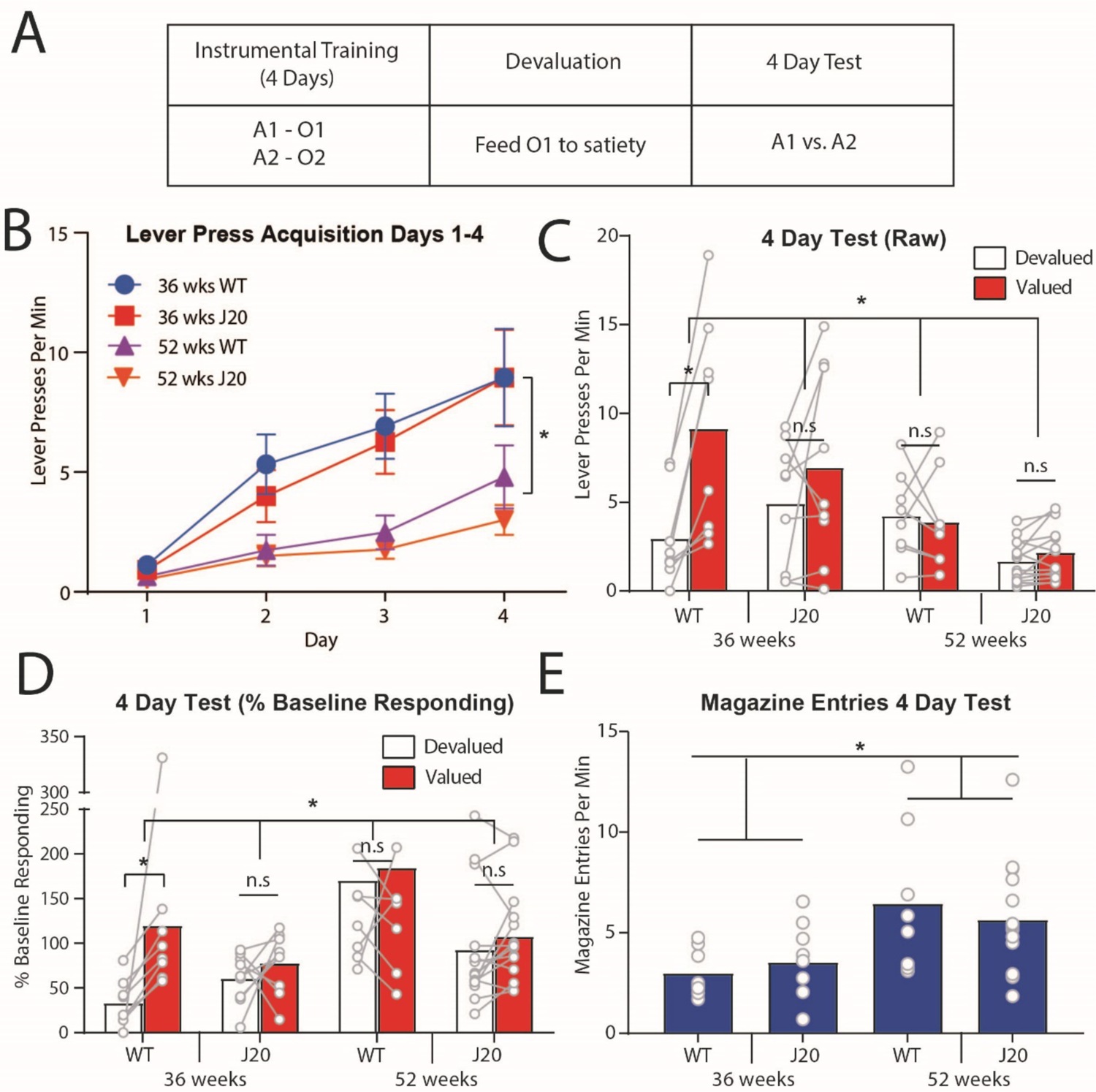
Outcome devaluation performance is initially impaired in 36-week-old male J20 mice and in all 52-week-old mice regardless of genotype. A) Design of outcome devaluation. B) Lever press rates (± SEM) during days 1-4 of lever press acquisition. C) Lever press rates during the 4 Day devaluation test. D) Lever press rates during the 4 Day devaluation test displayed as a percentage of baseline responding. E) Magazine entries during the 4 Day devaluation test. A = action, O = outcome.

Lever press rates during Days 1-4 of acquisition are shown in Figure 1B. Lever pressing increased across days for mice in all groups, although this increase was accelerated for the younger, 36-week-old mice relative to the older, 52-week-old mice. In support of these observations, there was a linear main effect, F(1,35) = 72.628, p = .00, and a main effect of age, F (1, 35) = 15.694, p = .00, but no main effect of genotype, F < 1. Importantly, all linear simple effects were significant (smallest F(1,35) = 4.978, p = .032, for 52-week-old J20s) demonstrating that mice in each group did increase their lever pressing across days, indicating intact lever press acquisition despite the overall lower press rates in 52-week-olds.

Lever pressing during the 4 Day devaluation test is shown in Figure 1C. As expected, devaluation was intact (Valued > Devalued) for 36-week-old wildtypes, and impaired (Valued = Devalued) for J20 mice of both ages. Unexpectedly, however, devaluation performance was also impaired for 52-week-old wildtype mice. Statistically, there was a significant 3-way (genotype x age x devaluation) interaction, F(1,35) = 5.059, p = .031, suggesting that devaluation performance differed between J20 mice and wildtype mice at 36-weeks-old, but not at 52-weeks-old. This interaction can be explained by the significant two-way genotype x devaluation interaction at 36-weeks, F(1,35) = 6.399, p = .016, but no such interaction at 52-weeks, F < 1. Specifically, at 36-weeks-old devaluation was intact (Valued > Devalued) for wildtypes, simple effect F(1,35) = 26.913, p = .000, but not J20s (Valued = Devalued) at this age, simple effect, F(1,35) = 3.294, p = .078, whereas devaluation was impaired and for both wildtype and J20 52-week-olds, both simple effects, Fs < 1.

Across this test, there was also a main effect of age F(1,35) = 9.821, p = .003 (but not of genotype, F(1,35) = 1.369, p = .25) suggesting that the younger mice, regardless of genotype, displayed higher overall levels of lever pressing (i.e., averaged across both levers). This finding, along with the main effect of age observed during lever press acquisition, raises the possibility that the older mice simply experienced a general motor impairment, as opposed to a specific impairment in their ability to exert goal-directed control. This conclusion is challenged, however, by the fact that 52-week-old mice made significantly *more* of the competing motor responses: head entries into the food magazine (Figure 1E), relative to 36-week-old mice averaged over genotype: main effect of age, F(1,35) = 10.416, p = .003, no main effect of genotype and no genotype x age interaction, both Fs < 1, suggesting that overall locomotor activity was equivalent in both groups.

Moreover, when lever press data from the test is calculated as a percentage of baseline responding, as shown in Figure 1D, this again produces a main effect of age, F(1,35) = 6.488, p = .015, but this is actually in the opposite direction, with 52-week-old mice responding *more* than 36-week-old mice relative to baseline. Yet the distribution of their responding remains equal between the valued and devalued levers. This is supported by a significant 2-way genotype x devaluation interaction for 36-week-old mice, F(1,35) = 4.608, p = .039, whereas there is no such interaction at 52-weeks, F < 1, and once again the 36-week-old wildtypes are the only group for whom a significant simple effect (Valued > Devalued) was observed, F(1,35) = 13.508, p = .001, whereas for all other group simple effects, F < 1. Together with magazine entry data, this result suggests that it is goal-directed control specifically, rather than motor control more generally, that is impaired in 52-week-old mice.

### Goal-directed action is intact for all male mice after additional lever press training, regardless of age or genotype

As mentioned, the devaluation impairment in J20 mice was expected to be transient. We tested this by giving mice an additional 4 days of lever press training (i.e., 8 days lever press training in total), and then administering a second devaluation test which we hereafter referred to as the “8 Day test”. On this test we expected devaluation performance to be intact for all mice, regardless of age or genotype.

Lever press rates during the additional lever press training (days 5-8) is shown in Figure 2A. Once again, 36-week-old mice lever pressed more than 52-week-olds, as supported by a main effect of age, F(1,35) = 14.049, p = .001, but lever pressing did not differ according to genotype, main effect, F < 1. Only 52-week-old mice increased lever press rates across days 5-8, as supported by a linear main effect, F(1,35) = 39.196, p = .00, that interacted with age, F(1,35) = 8.333, p = .007. Follow up simple effects reveal that this interaction comprises linear increases in both 52-week-old groups: wildtypes, F(1,35) = 32.913, p = .00 and J20s, F(1,35) = 13.436, p = .001, but no such increase for either group at 36 weeks of age: wildtypes, F(1,35) = 1.622, p = .211 and J20s, F(1,35) = 3.912, p = .056 (although the latter effect could be considered marginal).

**Figure 2:**
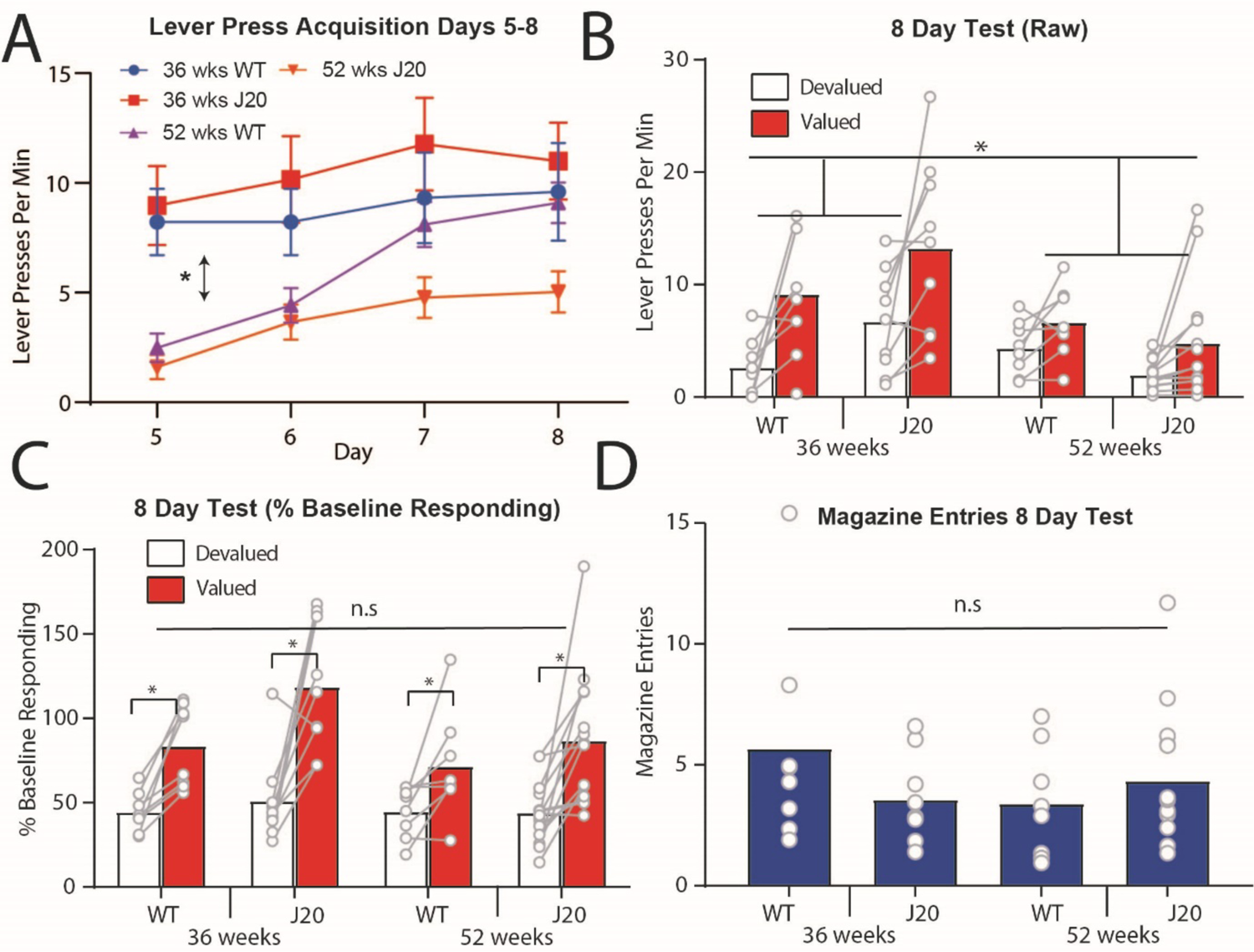
Outcome devaluation was intact in the 8 Day devaluation test for all male mice, regardless of age or genotype. A) Lever press rates (± SEM) during days 5-8 of lever press acquisition. B) Lever press rates during the 8 Day devaluation test. C) Lever press rates during the 8 Day devaluation test displayed as a percentage of baseline responding. D) Magazine entries during the 8 Day test.

Lever pressing during the 8 Day test is shown in Figure 2B. From this figure, devaluation performance appears intact (i.e., Valued > Devalued) for all groups despite the persistence of lower overall press rates in the 52-week-olds. These observations are supported by a main effect of age, F(1,35) = 9.152, p = .005, but not of genotype, F < 1, and a main effect of devaluation, F(1,35) = 27.95, p = .00. This time, however, there was no 3-way interaction, F < 1, and no two-way interactions at either 36-weeks or 52-weeks, both Fs < 1. There was, however, a two-way age x devaluation interaction, F(1, 35) = 5.527, p = .024, suggesting that performance was superior for 36-week-olds relative to 52-week-olds.

However, this result appears to be a consequence of the lower lever press rates in the older mice, rather than a difference in the ability to express goal-directed action control, because once test performance was expressed as a percentage of baseline responding these differences disappeared. These data are shown in Figure 2C. Specifically, there was again main effect of devaluation, F(1,35) = 49.607, p = .00, but this time there was no age x devaluation interaction, F(1,35) = 2.177, p = .149, suggesting that devaluation was indeed intact for all mice. Moreover, and in contrast to the 4 Day test, there were no group differences in magazine entries on this test as shown in Figure 2D: all Fs < 1.

Taken together with the results of the Day 4 test, these findings suggest that goal-directed action is initially impaired in J20 mice at 36-weeks old, and for mice of both genotypes at 52-weeks old, but that it is intact for all mice after additional lever press training.

### Outcome selective reinstatement is impaired for 52-week-old J20 males relative to wildtypes

So far, we have detected a wildtype/J20 difference in initial goal-directed control for 36-week-old mice but have failed to detect any genotypic differences for 52-week-olds. We therefore added another test of choice behaviour – outcome-selective reinstatement – in order to determine whether this lack of difference was general across tasks.

Following the 8 Day devaluation test, 52-week-old mice were re-trained on lever pressing for one day then subject to reinstatement testing. The design is shown in Figure 3A. For this test, animals were first exposed to 30 minutes of extinction on both levers. Mice then received two unsignalled presentations of each outcome in the following order: sucrose, pellets, pellets, sucrose, each separated by a further 4 minutes of extinction. If outcome-selective reinstatement were intact, animals should selectively reinstate responding on the lever that had earned that outcome during training. For instance, sucrose presentations should elicit pressing on the sucrose lever, and vice versa.

**Figure 3.**
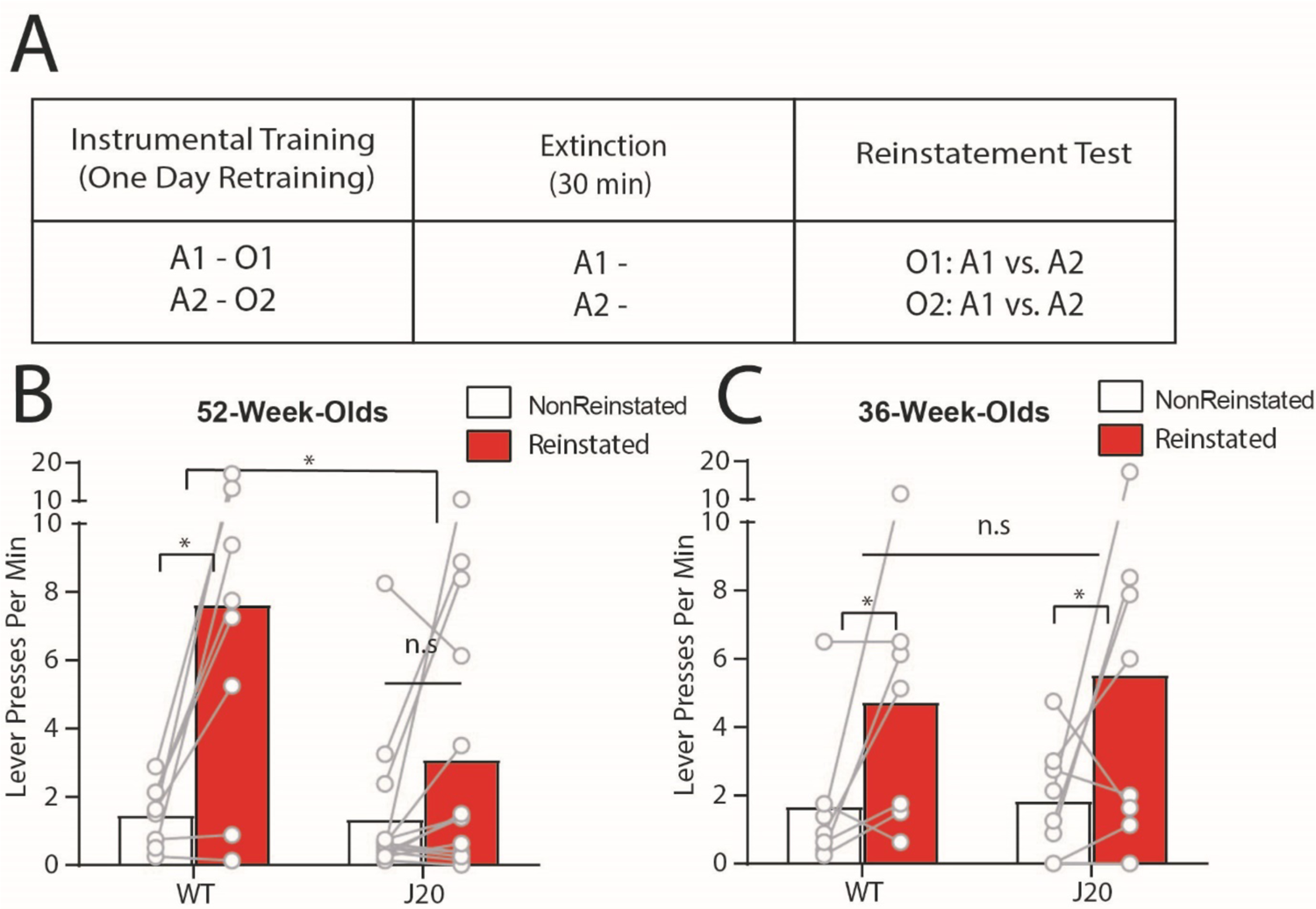
Outcome-selective reinstatement is impaired in 52-week-old J20 male mice but is unaltered in J20 male mice at 36-weeks-old. A) Design of the outcome selective reinstatement procedure t. B-C) Lever press rates during outcome selective reinstatement testing of B) 52 and C) 36-week-olds. A = action, O = outcome.

Test data is shown in Figure 3B. This time, we did observe a genotypic difference in 52-week-olds because selective reinstatement was intact for wildtypes (Reinstated > Nonreinstated) but impaired for J20s (Reinstated = Nonreinstated). Statistically, there was a main effect of reinstatement, F(1,20) 18.716, p = .00, no main effect of genotype, F(1,20) = 3.379, p = .081, and an age x reinstatement interaction, F(1,20) = 5.526, p = .029, supported by a significant simple effect for wildtypes, F(1,20) = 17.514, p = .00, but not J20s, F(1,20) = 2.683, p = .117.

Unfortunately, at the time the decision was made to test selective reinstatement, several of the 36-week-old male mice had already been culled. We thus decided to test reinstatement in an independent cohort of 36-week-old males (wildtypes: n = 7, J20s: n = 8), and confirmed that outcome selective reinstatement is intact for both wildtypes and J20s at this age, as shown in Figure 3C. Statistically, there was a main effect of reinstatement, F(1,13) = 6.626, p = .023, no main effect of genotype, and no genotype x reinstatement interaction, both Fs < 1.

This result demonstrates that one aspect of choice behaviour, selective reinstatement, is impaired in 52-week-old J20s relative to wildtypes. Together with the devaluation, this result suggests that after 8 days of training 52-week-old J20 mice are capable of using response-outcome associations, but not outcome-response associations to inform their action selection (Abiero et al., 2022; Balleine & Dickinson, 1996; Ostlund & Balleine, 2007).

### Goal-directed action is transiently impaired for 36-week-old J20 female mice

Nineteen females, comprised of n = 9 wildtypes and n = 10 J20s, were trained on a modified version of the outcome devaluation procedure previously described. Specifically, all procedures were identical except that the advancement through the ratio requirements for lever pressing was slower, and outcome devaluation tests were conducted for 5 minutes each rather than 10 minutes to reduce extinction (due to low levels of lever pressing, see materials and methods for details).

Lever press acquisition for females is shown in Figure 4A. Three mice were excluded from the analysis (2 WT and 1 J20) for failing to perform more than a single lever press during initial acquisition. Final sample size was therefore n = 7 wildtypes and n = 9 J20s. From Figure 4A lever pressing increased similarly for both groups across days as confirmed by a linear main effect, F(1,14) = 30.123, p = .00, no main effect of group, F(1,14) = 1.663, p = .218, and no group x linear interaction, F(1,14) = 1.511, p = .239.

**Figure 4.**
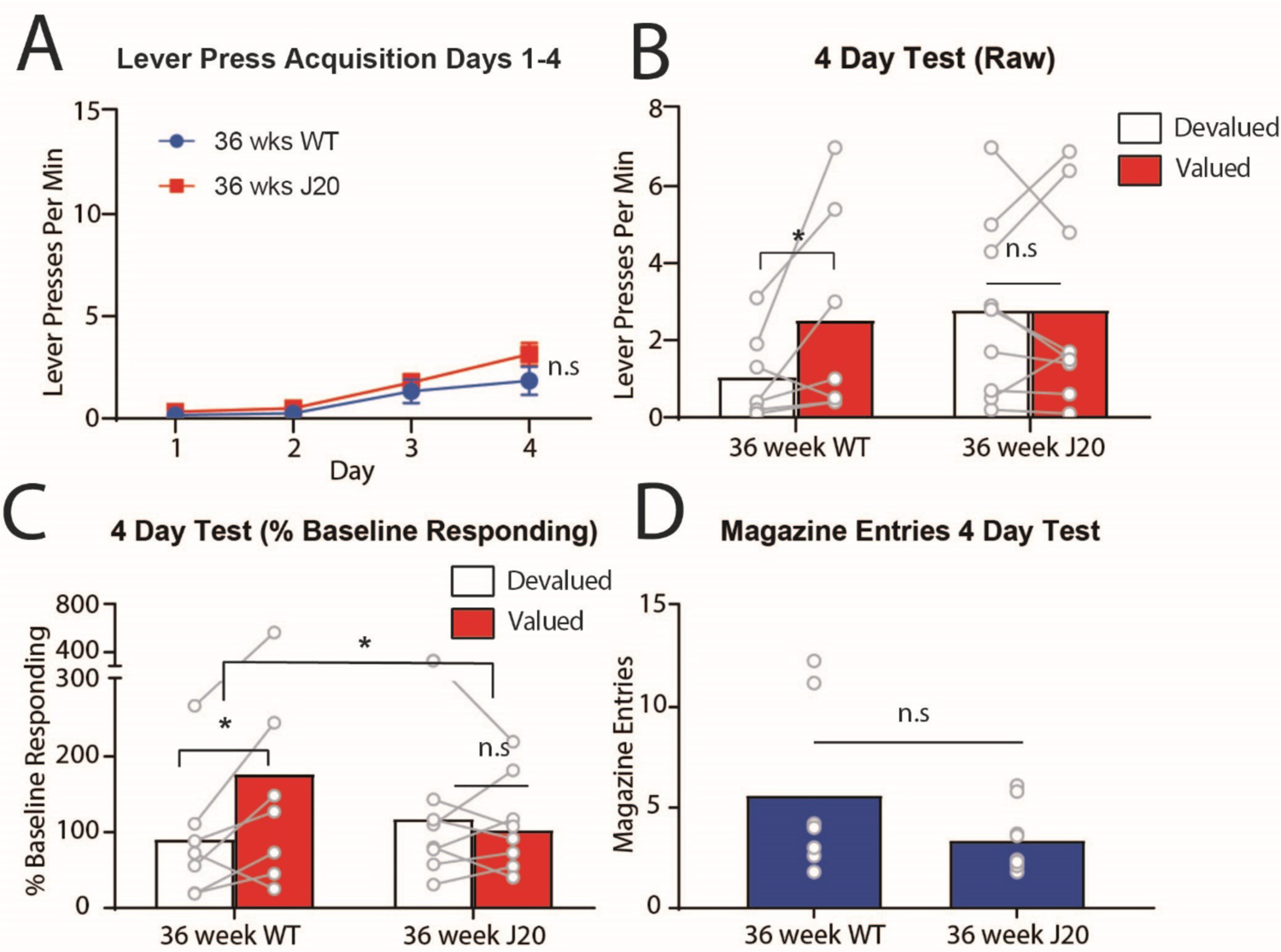
Outcome devaluation performance is initially impaired in female J20 mice at 36-weeks-old. A) Lever press rates (±SEM) during days 1-4 of acquisition. B) Lever press rates during the 4 Day devaluation test. C) Lever press rates during the 4 Day devaluation test displayed as a percentage of baseline responding. D) Magazine entries during the 4-day test.

Performance on the 4 Day devaluation test is shown in Figure 4B. This figure reveals a similar pattern of responding to that observed in the 36-week-old males: intact devaluation in wildtype mice (Valued > Devalued) and impaired devaluation in J20s (Valued = Devalued). Statistically, however, this result was not as robust as that observed for male mice (possibly due to lower sample size) as there was no main effect of group, F < 1, no main effect of devaluation, F(1,14) = 2.868, p = .112, and no group x devaluation interaction, F(1,14) = 2.868, p = .112. Nevertheless, once test data was transformed to a percentage of baseline responding, as shown in Figure 4C, although there was still no main effect of group, F < 1, and no main effect of devaluation, F(1,14) = 3.216, p = .095, there was a significant group x devaluation interaction, F(1,14) = 4.668, p .049. Follow-up analysis revealed a significant simple effect in the wildtypes, F(1,14) = 6.948, p = .02, suggesting that they pressed the valued lever more than the devalued lever, but no such simple effect for the J20s, F < 1, demonstrating that they pressed both levers equally. As shown in Figure 4E, magazine entries did not differ between groups on this test, F(1,14) = 2.084, p = .171.

Female mice were next subject to 4 more days of lever press training (i.e., 8 Days total) and retested for devaluation performance to determine if the initial impairment in J20 mice was transient.

Mice in both groups continued to acquire lever pressing across days 5-8 as shown in Figure 5A and supported by a linear main effect, F (1,14) = 108.45, p = .00, and this increase was equivalent because there was no main effect of group, F(1,14) = 2.271, p = .154, and no group x linear interaction, F(1,14) = 1.408, p = .255.

**Figure 5.**
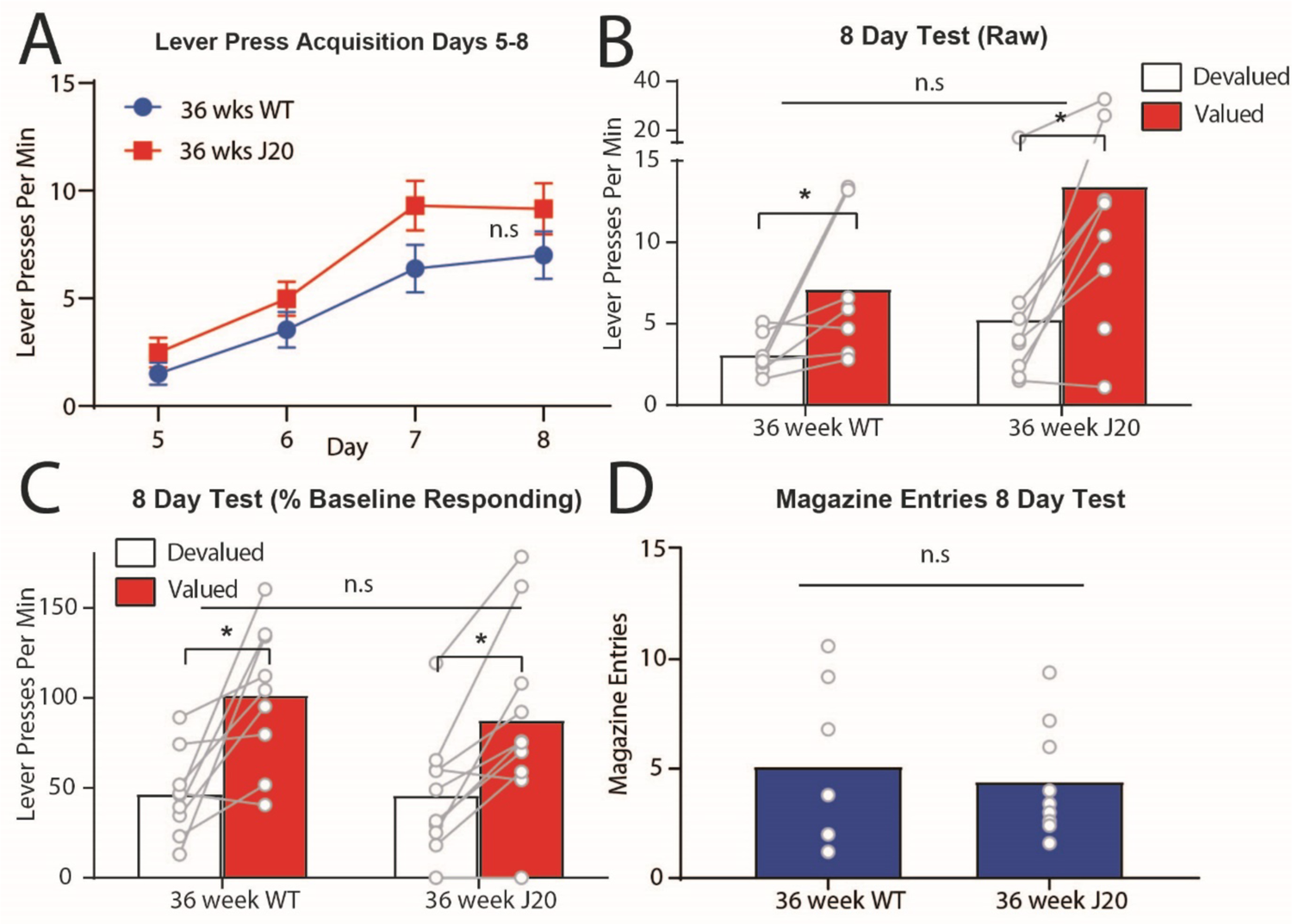
Outcome devaluation performance is intact for all female mice, regardless of genotype, after 8 days of lever press training. A) Lever press rates during days 5-8 of acquisition. B) Lever press rates during the 8 Day devaluation test. C) Lever press rates during the 8 Day devaluation test as a percentage of baseline responding. D) Number of entries into the food receptacle (magazine) averaged over days 5-8.

Data from the 8 Day test is shown in Figure 5B. It is clear from this figure that devaluation was intact (Valued > Devalued) for both groups, and this is supported by a main effect of devaluation, F(1,14) = 15.862, p = .001, no main effect of group, F(1,14) = 2.335, p = .149, and no group x devaluation interaction F(1,14) = 1.843, p = .196. Although not statistically supported, it does appear from Figure 5B as though mice in the J20 group were responding more overall than wildtypes on this test. However, if we again examine test performance as a percentage of baseline responding, as shown in Figure 5C, this difference dissipates. Specifically, there is a main effect of devaluation, F(1,14) = 30.582, p = .000, no main effect of group, F < 1, and no group x devaluation interaction F(1,14) = 1.129, p = .306. Magazine entries did not differ between groups on this test, F < 1, as shown in Figure 5D.

Together with data from the 4 Day test, these results suggest that goal-directed action is transiently impaired in female J20s relative to wildtypes, in a similar manner to the results for male mice of 36-weeks old. This suggests that the deficit is general to both sexes.

### Outcome-selective reinstatement is intact for all 36-week-old female mice

We also tested female mice for their performance on outcome-selective reinstatement, data from this test is shown in Figure 6. Outcome-selective reinstatement (Reinstated > Nonreinstated) was intact for all females, regardless of genotype. Specifically, there was a main effect of reinstatement, F(1, 14) = 10.49, p = 0.006, but no main effect of group, F(1,14) = 2.278, p = .153 and no group x reinstatement interaction, F < 1.

**Figure 6:**
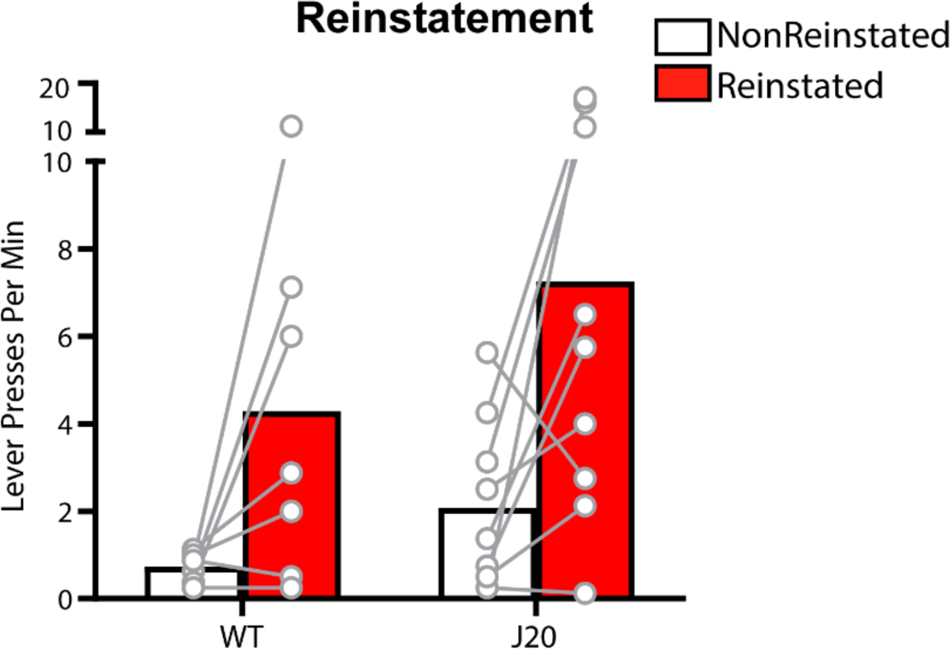
Outcome-selective reinstatement was intact for all females, regardless of genotype. Graph shows lever press rates for female 36-week-old mice during outcome selective reinstatement testing.

### Dorsal CA1 microglial (IBA1) expression increased with age but not genotype in male mice, and negatively correlated with initial devaluation performance

Following behavioural testing, all animals were perfused with 4% paraformaldehyde and brains were removing for sectioning and immunostaining with IBA1, a protein that is only present in microglia and is upregulated when those cells are activated (Ito et al., 1998) as occurs during an inflammatory response (Harry, 2013). Sections were co-stained with Amylo-Glo, to identify amyloid plaques (Schmued et al., 2012), see materials and methods for details. Representative examples of these stains from sections taken from wildtype and J20 mice at 36 and 52 weeks of age, respectively, are shown in Figure 7A-D. Sections were imaged were imaged and cell/plaque counts for each marker in CA1 region of the hippocampus at bregma −1.9mm. Due to a freezer malfunction, several of our brain samples were either lost or damaged to a point that we were unable to image them. Nevertheless, we were able to include the majority of samples in our analysis.

**Figure 7.**
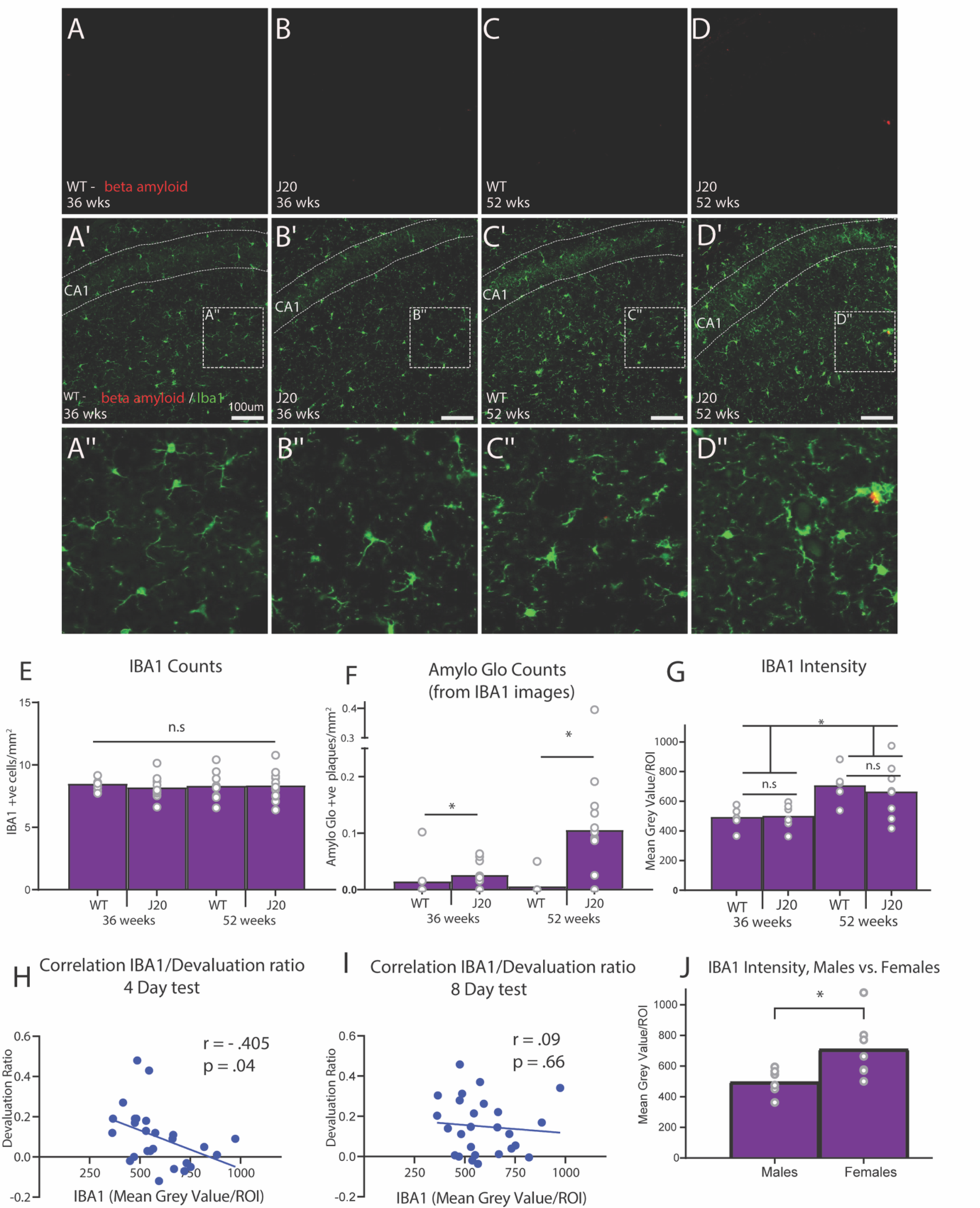
Quantification of IBA1 and Amylo Glo expression in the dorsal CA1 of male mice. A-D) Representative photomicrographs of male dorsal CA1 stained with Amylo Glo (red) in each group, scale bar is 100μm, A’-D’) Representative photomicrographs of male dorsal CA1 stained with Amylo Glo (red) and IBA1 (green) in each group, A’’-D’’) Enlarged inset, E) Number of cells positively immunostained for IBA1 per mm^2^, F) Number of plaques positively immunostained for Amylo Glo (taken from IBA1 images) per mm^2^, G) IBA1 intensity quantification, H) Correlation between IBA1 intensity and devaluation ratio from the 4 day test, I) Correlation between IBA1 intensity and devaluation ratio from the 8 day test, J) IBA1 intensity quantification for 36-week-old J20 males vs. 36-week-old J20 females.

First we analysed sections from male mice. Initially we analysed IBA1 and Amylo Glo expression by conducting automated counts using ImageJ software (see methods for details), but failed to find any group differences in IBA1 positive cells, Fs < 1, shown in Figure 7E. As expected, however, plaques stained for Amylo Glo were significantly higher in J20s than wildtypes (Figure 7F), there was a main effect of genotype, F = 6.318, p = .017, but not of age, F = 2.608, p = .116. Despite some obvious co-localisation between microglia and plaques, as can be seen in Figures 7A-C, the expression of IBA1 and Amylo-Glo for J20 did not significantly correlate, r = .047, p = .835. It should be noted that for the two WT mice for which ‘plaques’ were identified (Figure 7F) these were likely background as they did not colocalise with microglia as would be expected.

Following the failure to detect any group differences in microglia using cell counts, additional brain sections for each mouse were taken and immunostained for IBA1 and then imaged using a different, intensity-based quantification for which mean grey values were obtained for each marker (see materials and methods). Our reasoning was that although genotype and age may not lead to changes in number of cells, they could lead to alterations in cell morphology that might be detected by an intensity-based but not a count-based methodology. The results of this quantification are shown in Figure 7G. IBA1 expression did differ according to age using this analysis, but it still did not differ according to genotype, as confirmed by a main effect of age (52 weeks > 36 weeks), F(1,22) = 14.192, p = .001, but not genotype, F < 1, and no age x genotype interaction, F < 1.

We next correlated IBA1 expression with devaluation performance on both the 4 Day and the 8 Day tests. To calculate this correlation, we used a ‘devaluation score’ to ensure that any correlation detected wasn’t driven by baseline differences in lever press responding *per se.* For this score, we first calculated from suppression ratio (SR) scores on each of the levers (i.e., valued and devalued) according to the following equation:

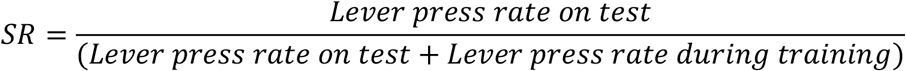

We then subtracted the Devalued SR from the Valued SR for each test separately, such that a positive devaluation score indicated more responding on the valued over the devalued lever, indicative of intact goal-directed action, and a zero or negative score indicated impaired goal-directed action. This score was correlated with microglial expression.

As shown in Figure H-I, we found that IBA1 expression (mean grey value) negatively correlated with devaluation ratio scores taken from the 4-day test, r = −.405, p = .04 (Figure 7H) but not the 8-day test, r = .09, p = .66 (Figure 7I). This result suggests that increased microglial expression in the dorsal CA1 is associated with poorer devaluation performance on the initial test but not after extended training, which could imply that initial deficits in goal-directed action are associated with neuroinflammation in this region. Amylo-Glo counts did not correlate with any behavioural measure on either test, smallest p = .157.

For female mice, we were only able to image and quantify sections from 3-4 wildtypes such that the sample size was not sufficiently large to allow an accurate statistical analysis of group differences. Furthermore, there were no significant correlations with these markers and any behavioural measures for female mice, although this was again likely due to low sample size. The quantification and correlation values for these mice are included in the data files at DOI 10.17605/OSF.IO/JXYC9, if researchers are interested in observing the numerical differences and/or conducting their own analyses. Nevertheless, because immunohistochemistry and subsequent quantification for the intensity-based analyses were conducted on all sections from male and female mice at the same time using the same techniques and parameters, we were able to directly compare levels of IBA1 between 36-week-old J20 animals of each sex (n = 7 females, n = 7 males) as shown in Figure 7J. IBA1 expression was significantly higher in female J20s relative to male J20 mice at this age, F(1,12) = 7.096, p = .021.

### Dorsal CA1 astrocytic (GFAP) expression in male mice increased marginally with age but not genotype, but did not correlate with any behavioural measure

Separate sections were taken from the same animals and co-stained with GFAP, a common marker for astrocytes, as well as with Amylo-Glo, to identify amyloid plaques (See materials and methods). Representative examples of these stains from wildtype and J20 mice at both 36 and 52-weeks of age, respectively, are shown in Figure 8A-D”. Sections were imaged and quantified as previously described. Analyses of cell counts for male mice are shown in Figures 8E-F. Again, counts of GFAP positive cells (Figure 8E) did not differ according to genotype, F < 1, or age, F(1,30) = 1.268, p = .269, whereas Amylo Glo positive plaques (Figure 8F) were significantly higher in J20s relative to wildtypes, F(1,30) = 8.058, p = .008, but not age, F < 1. Again, overall numbers of cells/plaques positive for GFAP and Amylo-Glo did not significantly correlate with each other, r = .12, p = .61. In addition, although ‘plaques’ were identified in two WT mice, these were also likely background staining.

**Figure 8.**
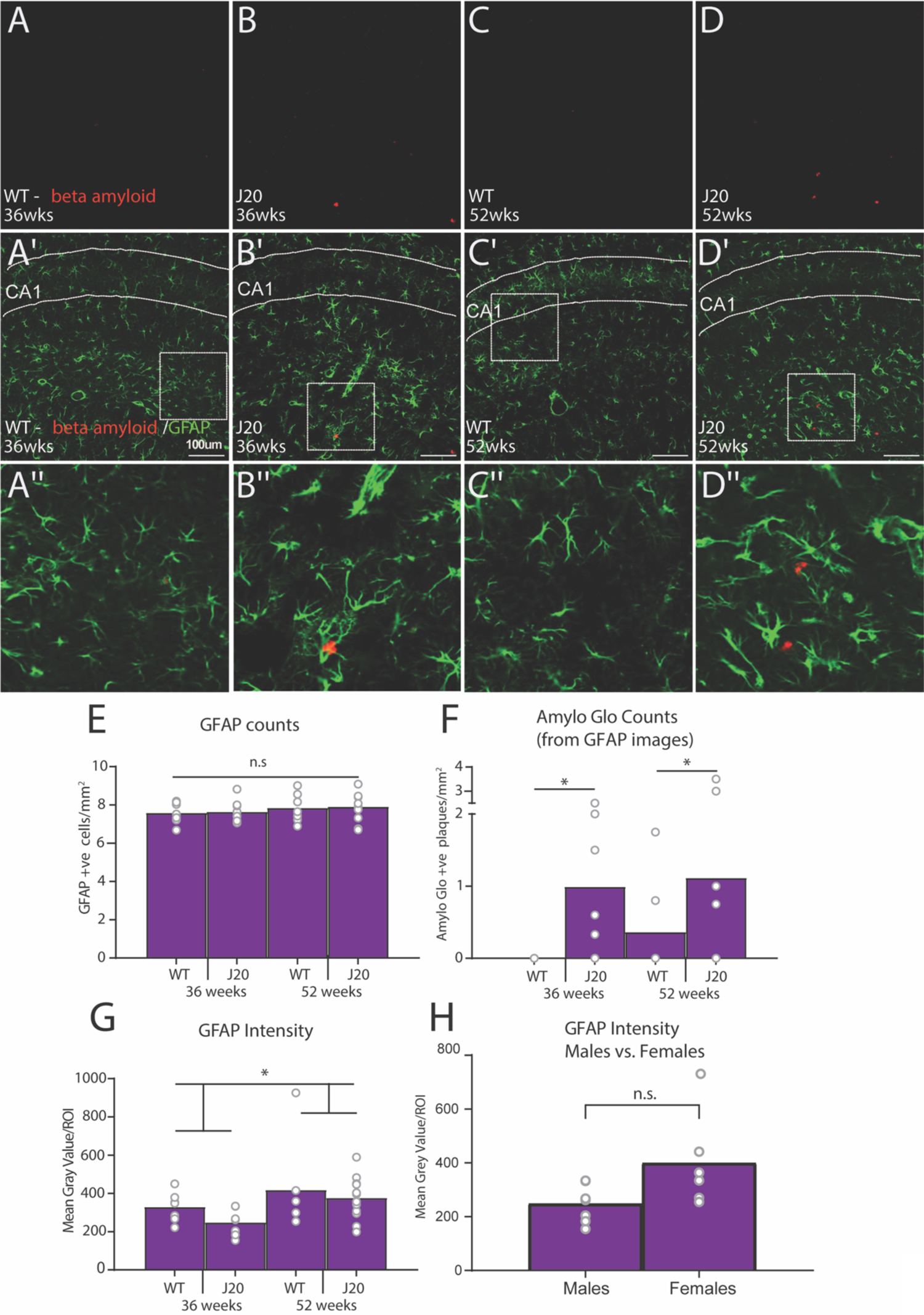
Quantification of GFAP and Amylo-Glo expression in the dorsal CA1 of male mice. A-D) Representative photomicrographs of male dorsal CA1 stained with Amylo Glo (red) in each group, scale bar is 100μm, A’-D’) Representative photomicrographs of male dorsal CA1 stained with Amylo Glo (red) and GFAP (green) in each group, A’’-D’’) Enlarged inset, E) Number of cells positively immunostained for GFAP per mm^2^, F) Number of plaques positively immunostained for Amylo-GLo (taken from GFAP images) per mm^2^, G) GFAP intensity quantification, H) GFAP intensity quantification, 36-week-old J20 males vs. 36-week-old J20 females.

Again, we conducted a separate immunostain and intensity-based analysis for GFAP with the hope of detecting morphology differences not obtained using cell counts. The results are shown in Figure 8G. From this figure, it appears that, like IBA1, GFAP expression did not differ according to genotype but that there was higher expression in the older, 52-week-old mice relative to 36-week-olds. Indeed, there was no main effect of genotype F(1,28) = 1.506, p = .23, but there was an effect of age, F(1,28) = 4.549, p = .042, that did not interact with genotype F < 1. GFAP expression did not significantly correlate with any behavioural measure on either test, all p’s > 0.05 (see data files for more detail).

Again, low sample size in wildtype females precluded an analysis of group differences of GFAP in the female cohort, however the quantification and correlation values for these mice are included in the data files at DOI 10.17605/OSF.IO/JXYC9. Once again, however, because immunohistochemistry and subsequent quantification for the intensity-based analyses were conducted on all sections from male and female mice at the same time for 36-week-old J20 animals of each sex (n = 7 males, n = 6 females), we were able to compare these directly as shown in Figure 8H. As with IBA1, GFAP expression was increased in female J20s relative to male J20 mice at this age, although statistically the effect was only marginal, F(1,11) = 4.39, p = .06.

## Discussion

Here we report several experiments demonstrating a transient impairment in goal-directed action in mice of both sexes from an hAPP-J20 preclinical mouse model of Alzheimer’s disease, and in aging male mice. Specifically, for both male and female 36-week-old mice, outcome devaluation was initially impaired for J20s (Valued = Devalued) relative to wildtypes (Valued > Devalued), whereas it was impaired for all 52-week-old male mice regardless of genotype. Following an additional 4 days of lever press training, outcome devaluation was intact for all mice regardless of age or genotype. On a separate test of choice behaviour: outcome-selective reinstatement, performance was intact for 36-week-old mice of both sexes and genotypes but was impaired for 52-week-old male J20s relative to age-matched wildtypes. Immunohistochemical analyses of dorsal CA1 hippocampal tissue revealed increased microglial expression (intensity) in 52-week-old males relative to 36-week-old males, regardless of genotype, and this expression negatively correlated with devaluation performance on the 4 Day, but not the 8 Day, test. Finally, our immunohistochemical markers of microglia (IBA1) and astrocytes (GFAP) were increased in 36-week-old female J20s relative to male J20s of the same age, a result that appears to mirror the enhanced prevalence of Alzheimer’s disease in female patients relative to male patients, for whom it is approximately twice as frequent (Podcasy & Epperson, 2016; Laws, Irvine, & Gale, 2018). Together, these results suggest that goal-directed action and choice behaviour is altered in an hAPP-J20 preclinical model of Alzheimer’s disease, and that the impairments are associated with dorsal hippocampal neuroinflammation.

It is first worth addressing the puzzling result that our putative neuroinflammatory markers (IBA1 and GFAP) did not differ according to genotype. Specifically, we had expected to see J20 > wildtype differences in these markers, as has been previously observed (Flores et al., 2018; 2021). There are several things to note here. First, our quantification techniques could be seen as less thorough than the stereological techniques used by some prior researchers (e.g. Wright et al., 2013), which could pick up smaller or more nuanced differences in the data. Second, IBA1 is a marker of all microglia, and it may be that only microglia that become activated as part of a (presumed) neuroinflammatory response differ according to the J20 genotype (e.g. Hong et al., 2016; Wright et al., 2013). Third, GFAP is not an absolute marker of all non-reactive astrocytes, and although its immunoreactivity is quite strong in the hippocampus as compared to other markers, it doesn’t capture the entire astrocyte population, as well as the complete morphology of the cell (Jurga et al., 2021; Zhang et al., 2019). Finally, other studies report findings that are consistent with current results, having failed to find any such differences in the dorsal CA1 region specifically (Dekens et al., 2018; Wright et al., 2013) or between J20s and age-matched wildtypes at 52-weeks-old (Ameen-Ali et al., 2019), suggesting that any genotypic differences observed with these markers could be highly region-and age-specific.

The behavioural finding that goal-directed action was transiently impaired in 36-week-old mice of both sexes parallels previous findings of a transient role for the dorsal CA1 region of the hippocampus in goal-directed action (Bradfield et al., 2020). To our knowledge, this is the only brain region that has been identified upon which goal-directed action relies initially, but not after additional lever press training. Lesions and/or inactivations of alternate brain regions involved in goal-directed action, such as posterior dorsomedial striatum (Yin et al., 2005), nucleus accumbens core (Corbit et al., 2001), prelimbic cortex (Corbit & Balleine, 2003), basolateral amygdala (Ostlund & Balleine, 2008), medial orbitofrontal cortex (Bradfield et al., 2015; 2018), and mediodorsal thalamus (Corbit et al., 2003), have all produced impairments in goal-directed action that are sustained even after multiple days of lever press training (although we note that this is not always the case if the manipulation occurs post-training). Thus, the pattern of behavioural results in the current study, coupled with the negative correlation between dorsal CA1 microglial expression and devaluation performance, strongly suggests that the goal-directed deficits observed here were also a result of damage to this particular region.

Current findings produced the additional surprising result that goal-directed action is transiently impaired as a result of aging. That is, devaluation performance was initially impaired for both wildtype and J20 mice at 52-weeks-old. Correspondingly, microglial (and to a lesser extent, astrocytic) expression also appeared to increase with age, being higher in 52-week-old male mice relative to their 36-week-old counterparts. Although behavioural impairments and microglial expression have both previously been found to increase with age (e.g., Matamales et al, 2016; Schilling & Eder, 2015), what stands out about current results is the relatively young age at which these developments occurred. That is, almost all prior studies have identified such differences comparing younger mice to mice aged 18 months or older – an age that is at least 6 months older than our oldest mice. If our current results are replicable and translatable, this finding could suggest that changes to brain and behaviour might occur earlier than previously thought.

As noted, devaluation performance did not differ between J20 and wildtype mice at 52-weeks-old, but we did identify an impairment in the performance of another type of choice behaviour that was specific to 52-week-old J20s: outcome-selective reinstatement. This result is interesting for two reasons. First, it highlights the changing nature of the cognitive-behavioural deficits of J20 mice as they age, suggesting that functions that are intact for younger J20 mice later become impaired, mirroring the cognitive decline that occurs in humans as Alzheimer’s disease progresses. Second, it suggests that the neuroanatomical location of the impairment might also have spread throughout the brain because we have recently presented evidence (Abiero et al., 2022) that outcome-selective reinstatement performance is unlikely to rely on dorsal hippocampus. Nevertheless, it isn’t immediately clear what brain region could be mediating this particular effect because, to our knowledge, no brain region has been identified to mediate selective reinstatement but not outcome devaluation (which was intact in these animals on the 8 Day test). Thus, this question will remain to be answered by future studies. Regardless, this finding does suggest that selective reinstatement could prove to be a particularly important test for detecting differences between older wildtype mice and transgenic mice engineered to mimic aspects of Alzheimer’s disease.

From a cognitive perspective, we have previously suggested that the transience of the dorsal hippocampus’ role in goal-directed action is reflective of this region’s role in episodic memory (Bradfield et al., 2020; Abiero & Bradfield, 2021). Specifically, we have suggested that goal-directed action initially relies on episodic memory but becomes dependent on extra-hippocampally encoded semantic memory over time. Such a theory fits neatly with current results because episodic memory impairments are well-known early indicators of Alzheimer’s disease (Rémy et al., 2005; Gold & Budson, 2008) with semantic deficits appearing only as the disease progresses (Sánchez et al., 2017). That is, it is possible that J20s and in 52-week-old male wildtypes suffered impaired episodic-like memory which led to their initial deficit in goal-directed action, but that intact semantic-like memories in all mice allowed for intact goal-directed performance on the 8 Day test. Such an interpretation is, of course, highly speculative, but does nicely illustrate the intersection of memory, motor control, and action selection as required for diagnoses of Alzheimer’s disease in humans.

Finally, the transience of the deficit observed here also has some important potential translational implications. Specifically, these results suggest that additional training allowed J20 mice to overcome their impairment to demonstrate intact goal-directed action. If translatable, this could suggest that individuals with Alzheimer’s disease might be similarly impaired when initially learning how to perform a new action or when learning to perform an action in a new environment. However, with enough training, repetition, and/or instruction, they could overcome this impairment to perform the action appropriately. Indeed, such behavioural interventions are reminiscent of the ‘dementia villages’ that have begun to be built around the world (Peoples, Pedersen & Moestrup, 2020). These villages allow individuals more time and help to make decisions about, for example, crossing the road, or what they would like to purchase from the shops. Our study provides additional validation for these kinds of interventions, even suggesting that they can even be modelled to an extent in mice.

Overall, we hope that the findings reported here provide an important starting point for researchers wishing to test a more comprehensive set of behavioural assays in preclinical mouse models of Alzheimer’s disease. Although spatial and contextual fear memory tests have been routinely included in the set of behavioural assays used to test rodent models of Alzheimer’s disease, the current study is the first, to our knowledge, that investigates how such memory deficits might translate into poorer cognitive control. In parallel, individuals with Alzheimer’s disease will only reach a diagnosis if their memory deficits impair their ability to function independently. Therefore, we hope that these findings will replicate in distinct mouse models of Alzheimer’s disease, and to this end, we are more than happy to provide additional details on our protocols or any other aspect of our design that we have neglected to mention here. We have also shared all of our data online at the following DOI: 10.17605/OSF.IO/JXYC9, should researchers want to conduct their own analyses of it.

## Materials and Methods

A total of 58 hemizygous transgenic (hAPP-J20) and non-transgenic mice (WT) were used from the J20 line, which express hAPP containing both the Swedish and Indiana mutations, under a PDGF-β chain promoter. Four mice were excluded for failing to learn to lever press (i.e., for performing no more than a single lever press prior to the 4 Day Test), and 1 was found dead, possibly as a result of a seizure which this mouse line is prone to (Palop et al., 2007). Thirty-nine of the remaining mice were males, 15 of which were approximately 36-weeks at the start of behavioural testing and 24 were approximately 52-weeks. All females were approximately 36-weeks at the start of testing as no older females were available. Mice were housed in sibling groups of 2-5, separated by sex.

Mice were bred and housed at Australian BioResources located in Mossvale, New South Wales, Australia until their issue to Garvan Institute, New South Wales, Australia. As litters reached the desired age range at different times, behavioural testing occurred in a staggered fashion (i.e., the age at which animals in the same group were in the age range for testing did not always overlap). However, all experimental and non-experimental (e.g. housing) variables were kept as consistent as possible to minimize any potential impacts of variables other than the target variables influencing performance. Upon arrival, mice weighed between 15-25g (females) and 20-40g (males).

All mice were maintained at a 12-hour 7am-7pm light/dark cycle and all experimentation was conducted in the light portion of the cycle. Mice were given *ad libitum* access to food and water until the experimentation began. Three days prior to behavioural training, the mice underwent dietary restrictions whereby each mouse was given 1-2 grams of chow per day, and unlimited access to water. These food restrictions were upheld for the duration of the experiment. During this time, mice were handled and weighed every second day to maintain their weight at >80% of their baseline body weight. All animal experimentations were performed with the approval of the Garvan Institute Animal Ethics Committee, in accordance with the National Health and Medical Research Council animal experimentation guidelines and Australian Code of Practice for the Care and Use of Animals for Scientific Purposes (2013).

### Apparatus

The training and experimentation were conducted in six identical operant chambers (Med Associates; St. Albans, VT, USA) equipped with a pump that delivered a 20% sucrose solution and a pellet dispenser that delivered a single grain of pellet (Able scientific; Perth, Australia) into the magazine (i.e. food receptacle) located in the middle of the side wall. The chamber also came equipped with two retractable levers, placed on either side of the magazine, a house light for illumination that was situated opposite the magazine, and a house fan which provided constant ~ 70db background noise. MED-PC software controlled the insertion of levers, delivery of pellet and sucrose outcomes, and recorded the number of lever presses and magazine entries.

### Behavioural procedures

#### Magazine training

Mice first received two sessions of magazine training over two days. The start of the session was signalled by illumination of the house light. During the session, sucrose and pellet outcomes were delivered to the magazine at random intervals around a mean of 60 seconds (i.e. on a random time 60 schedule). The session terminated after 30 minutes or after 20 of each outcome (40 outcomes in total) had been delivered, whichever came first. Levers were not extended during magazine training.

#### Lever press training, Days 1-4

One day after magazine training finished, mice were trained to lever press for 4 days. Each lever press training session lasted for 50 minutes and consisted of two 10 minutes periods on each lever (i.e., 4 x 10 minutes sessions in total) separated by 2.5 min time-out period in which the levers were retracted, and the house light switched off. Lever press periods terminated early if 20 outcomes were earned such that mice could earn a maximum of 40 pellets and 40 deliveries of sucrose solution per session. Contingencies were counterbalanced so that half the animals in each group received left lever-pellets, right lever sucrose, and the remaining half received the opposite arrangement.

For males, the first day of lever press training was continually reinforced (CRF) (i.e. each lever press was rewarded with an outcome). They were then shifted to a random ratio (RR-) 5 schedule for the next 2 days (i.e., each lever earned an outcome with a probability of 0.2), then to a RR-10 schedule (i.e., each lever earned an outcome with a probability of 0.1). Female mice were trained on CRF schedules for two days, then moved onto one day of RR5 and then one day of RR10. There was some variability according to individual press rates (i.e. slower pressers were kept on richer reward schedules for longer).

#### Outcome devaluation, 4 Day test

The first round of devaluation testing that occurred after 4 days of lever press training is referred to as the “4 Day Test”. For these tests, mice were each placed in an empty Perspex vivarium box and were given *ad libitum* access to either pellets or sucrose (counterbalanced) for one hour to induce specific satiety, reducing its value relative to the other outcome (Balleine & Dickinson, 1998). Immediately following devaluation, male mice were placed back in the operant chambers for 10 minutes, whereas female mice were placed in the operant chambers for 5 minutes. Both the levers were extended, but neither outcome was actually delivered (i.e., test was conducted in extinction). On day 2 of devaluation testing, 24 hours after the first, animals were pre-fed the alternative food source (i.e., if they were pre-fed pellets on day 1 they received sucrose, and vice versa) and were again tested in extinction for 10 minutes (males) or 5 minutes (females). Test results are reported as averaged across these two days of testing.

#### Lever Press training, Days 5-8

One day after the 4 Day devaluation test, all mice received 4 more days of lever press training. These training sessions took place as described above. All mice, regardless of sex, were trained on CRF, RR5, RR10, RR10 over days 5-8.

#### Outcome devaluation, 8 Day test

Mice were again subject to outcome devaluation testing after 8 total days of lever press training, in what we referred to as the “8 Day Test”. This test was conducted identically to the 4 Day test, described above.

#### Outcome-selective reinstatement

Outcome-selective reinstatement sessions began with 30 minutes of extinction, during which the house light was turned on and both levers extended and lever presses recorded, but no outcomes were delivered. During reinstatement, mice received 4 reinstatement trials separated by 4 minutes each. Each reinstatement trial consisted of a single free delivery of either the sucrose solution or the grain pellet. All rats received the same trial order: sucrose, pellet, pellet, sucrose. Responding was measured during the 2-minute periods immediately before (Pre) and after (Post) each delivery.

### Perfusions

After behavioural testing was finished, each mouse was injected intraperitoneally with a combination of ketamine (2mg/mL) and xylazine (8mg/mL), according to their individual weight to deeply anesthetize them. Once anesthetized, the mice were cut open from their abdominal region until the incision reached the ribcage, and the heart was exposed. The apex of the heart was punctured with a needle and a tiny incision was made on the right atrium, allowing the excess liquid to drain out. Firstly, the needle delivered a saline solution which pumped through the blood vessels of the mice to flush out the blood. Following this, the mice were transcardially perfused with 4% paraformaldehyde (PFA). Next, the brains were harvested and postfixed for 24 hours in 4% PFA in order to prevent it from decaying by terminating any biological reactions. The brains were then transferred to a 30% sucrose solution and finally sectioned coronally (40 μm) using a cryostat (Leica Biosystems, Switzerland). The sectioned slices were immediately immersed in cryoprotectant solution and stored at −20°C.

### Immunohistochemistry

Five representative anterior-posterior sections (coordinates of the sections ranged from Bregma −1.90 mm to Bregma −2.2 mm) from the CA1 region of the dorsal hippocampus were selected for each mouse for IBA1/Amylo-Glo staining procedures to stain for microglia and amyloid plaques, respectively. Additionally, 5 separate sections from the same region were selected for staining with both GFAP and Amylo-Glo to detect astrocyte expression and amyloid plaques respectively. All sections were first washed three times (10 minutes per wash) in sterile PBS (pH 7.2) to remove any exogenous substances. The sections were then incubated in a blocking solution comprising of 3% Bovine Serum Albumin (BSA) +0.25% TritonX-100 in 1x PBS for one hour to permeabilize tissue and block any non-specific binding.

### Ionized calcium binding adaptor molecule (IBA1)

Following permeabilization treatments and 3 more 10-minute washes in PBS, sections were first incubated in the primary antibody (1:1000 rabbit-IBA1, Abcam, Sydney, Australia) diluted in the same blocking solution described above for 72 hours at 4°C. Next, the sections were washed 3 more times in PBS followed by incubation in the secondary antibody (antirabbit Alexa Fluor-488, Abcam; Sydney, Australia) diluted in blocking solution (1:250) and preserved overnight at 4°C. The sections were washed in PBS for a final time (3x 10 minutes) before being stained by Amylo-Glo as per the procedure detailed below.

### Glial fibrillary acidic protein (GFAP)

The procedure for GFAP staining was identical to that described above as above for IBA1 except that the primary antibody was anti-GFAP (goat-GFAP, Abcam; Sydney, Australia) diluted at 1:300, and the secondary antibody was Alexa Fluor-488 (anti-goat Alexa-Fluor 488, Abcam; Sydney, Australia) diluted at 1:1000. The sections were washed in PBS for a final time (3x 10 mins) before being stained by Amylo-Glo.

### Amylo-Glo

Amylo-Glo was used to quantify the number of amyloid plaques in the brain and was used by diluting at 1:500 with 0.9% saline solution. After the sections were stained for IBA1 and GFAP, they were put into the Amylo-Glo solution for an hour. Sections were then mounted on SuperFrost-plus slides and cover slipped with Vectashield.

### Cell count quantifications of microglia, astrocytes and amyloid plaques

Selected CA1 sections that were immuno-stained for IBA1 and Amylo Glo, or GFAP and Amylo Glo using the same methods described previously were analysed under an Axio Imager.Z2 fluorescent microscope (Zeiss; Jena, Germany), and images of the CA1 region of the hippocampus of both right and left hemispheres were obtained. These images were then analysed and cells or plaques counted on ImageJ as follows: The image was adjusted to 8-bit and the background subtracted. The threshold for contrast and brightness was adjusted for all images until consistent between images. The size of the cells that ImageJ identified as IBA1 were set between 60-600 and set at 80-2000 for GFAP. The integrated density of those cells was measured.

ImageJ counted each cell between our parameters and presented it as a ‘count’. The area of the CA1 being counted was 19.6mm^2^, hence, these raw counts were divided by 19.6 to give us counts per mm^2^.

### Intensity quantifications of microglia, astrocytes and amyloid plaques

The stained and mounted sections were analysed under a confocal laser scanning microscope (Zeiss-LSM-7110 CLSM, Carl Zeiss, Germany) with a 20x optical magnification, resolution 1024 x 1024px and a Kaplan filter (3 average scans), centred in the acquisition area. Laser intensity, PMT voltage and offset were maintained constant in all acquisition of the same double immunofluorescence experiment. Images of the region of interest (ROI) of 635.9 x 635.9 mm of the CA1 region of in the hippocampus around bregma −1.9mm (Paxinos and Franklin, 2001) (of both left and right hemispheres) were obtained. R. aw 16-bit images were then analysed using ImageJ software (MacBiophotonics upgrade v. 1.43u, Wayne Rasband, National Institutes of Health, USA). For each marker a mean grey value was obtained from two to five sections and averaged for each animal. All measurements were quantified by a researcher who was blind to the group identity. The representative images were chosen and imported to ImageJ where the double stained IBA1/Amylo-Glo or GFAP/ Amylo-Glo were counted. An average of 5 sections were counted.

Amyloid plaque quantification was performed as Song et al., (2021). The captured images were analysed for Amylo-Glo intensity in ImageJ Fiji 2.0.0 (https://imagej.net/Fiji). We adjusted the images to 8-bit and subtracted the background. We then selected the proper threshold of signals (maximum: 255, minimum: 0). When all the Amylo-Glo signals in the section has become red, we quantified the Integrated density (total intensity within threshold). The Amylo-Glo intensity was measured in both hemispheres in two to five sections and were averaged for each mouse. All measurements were quantified by a researcher who was blind to the group identity of the sample.

## Acknowledgements

This work was funded by GNT2003346 from the National Health and Medical Research Council of Australia awarded to L.A.B, and DP200102445 from the Australian Research Council awarded to L.A.B. and B.V.

